# Specificity and overlap in the genetic architectures of functional and structural connectivity within cerebral resting-state networks

**DOI:** 10.1101/2022.05.31.494125

**Authors:** E.P. Tissink, J. Werme, S.C. de Lange, J.E. Savage, Y. Wei, C.A. de Leeuw, M. Nagel, D. Posthuma, M.P. van den Heuvel

## Abstract

The functional connectivity and dynamics of resting-state networks (RSN-FC) are vital for cognitive functioning. RSN-FC is heritable and partially translates to the anatomical architecture of white matter, but the genetic component of structural connections of RSNs (RSN-SC) and their potential genetic overlap with RSN-FC remains unknown. Here we perform genome-wide association studies (N_discovery_=24,336; N_replication_=3,412) and in silico annotation on RSN-SC and RSN-FC. We identify the first genes for visual network-SC, that are involved in axon guidance and synaptic functioning and show that genetic variation in RSN-FC impacts biological processes related to brain disorders that have previously been associated with FC alterations in those same RSNs. Correlations of the genetic components of RSNs are mostly observed within the functional domain, whereas less overlap is observed within the structural domain and between the functional and structural domains. This study advances the understanding of the complex functional organization of the brain and its structural underpinnings from a genetics viewpoint.

## Introduction

Structural (SC) and functional connectivity (FC) are vital for healthy cognitive behaviour^1^. Brain regions that show temporally synchronized activity form functionally specialized resting-state networks (RSNs)^2^, including primary networks (such as the visual or somatomotor network) and higher-order cognitive networks (such as the frontoparietal network, salience network, or default mode network)^3^. Many psychiatric and neurological disorders have been associated with disruptions within specific RSNs^4^ and improving our understanding of the biological principles underlying the concept SC and FC of RSNs (RSN-SC/FC) could help elucidate the neural basis of human cognition and disorders associated with disruptions in brain connectivity.

Studies have shown that genetic factors significantly contribute to RSN-FC (*H^2^* = 20-40%)^5–10^. Genome-wide association studies (GWAS) on FC graph theory measures^11^ and extrinsic and intrinsic functional organization^12^ of RSNs have identified the first genetic variants and genes that make up this genetic component (mean 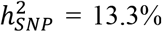^11^), and show genetic overlap between FC and psychiatric disorders^13^. RSNs were traditionally discovered based on FC^2^ and correlate with the structural connectivity (SC) architecture of white matter in the brain^14–16^ to varying degrees across RSNs^17^. The genetic architecture of RSNs-SC has not been investigated to date, but the substantial heritability of multiple properties of major white matter tracts (mean 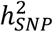 25.18% - 34.9%)^18–20^ suggests the importance of genetic factors for the anatomical backbone of RSNs. Describing the genetic architecture of both RSN-FC and RSN-SC as well as annotation and interpretation of the genetic signal can give insight into a biological substrate relevant to a wide variety of neurological and psychiatric disorders^21^ and additionally enables us to estimate to which degree RSN-SC relates to RSN-FC based on a shared genetic source.

In this study, we aim to characterise the genetic architecture of RSNs, both structurally and functionally. Large-scale (discovery N_FC_ = 24,336 and N_SC_ = 23,985; replication N_FC_ = 3,408 and N_SC_ = 3,412) GWAS are performed on the SC and FC within seven well-known RSNs^2^. We estimate and partition the SNP-based heritability and examine the convergence of the polygenic signal from these GWAS onto genes and biological pathways, with the purpose of aiding the biological interpretation of results and providing meaningful starting points for functional follow-up experiments^22^. We examine genetic correlations both between different RSNs, as well as across structural and functional domains. These genetic correlation analyses are extended to the locus level to facilitate the prioritisation of possible pleiotropic loci for future studies^23^. Altogether, we focus on the translation of RSN-associated genetic loci into biological interpretation and provide insights into the genetic specificity and overlap of RSN-FC and RSN-SC.

## Results

### GWAS of RSN-SC and RSN-FC identify six genome-wide significant loci

Following previously described procedures^24^, we started our analysis by grouping cortical areas into seven RSN as defined by Yeo et al^2^ (visual, somatomotor, limbic, dorsal attention, ventral attention, frontoparietal, and default-mode network; Supplementary Figure 1) and calculating the mean functional and structural connectivity within the RSNs in UK Biobank subjects (discovery N_FC_ = 24,336 and N_SC_ = 23,985; replication N_FC_ = 3,408 and N_SC_ = 3,412). RSN functional connectivity was measured as the average correlation between the activation signals of brain regions within each RSN over time, RSN structural connectivity was measured as the average fractional anisotropy (FA) of white matter tracts between brain regions within each RSN (see Methods). Discovery GWAS were performed for the FC and SC within every RSN and identified 518 genome-wide significant SNPs (*p* < 5×10^-8^/16 = 3.13×10^-9^) located in six genomic loci: three for visual network-SC, one for limbic network-FC, and a shared locus for frontoparietal network-FC and somatomotor network-FC (Supplementary Table 1). These loci seem to show RSN specific genetic effects rather than simply being driven by overall connectivity, given that none of these six loci showed a genome-wide significant association with global FC or SC.

SNP-based heritability 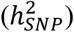 estimates for RSN-SC (M = 13.59%, SD = 1.79%) were moderately higher than those observed for RSN-FC (M = 6.71%, SD = 3.36%; Supplementary Table 2). We did not find evidence for enrichments of 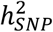 in functional genomic categories after Bonferroni-correction (Supplementary Methods 1.1 and Supplementary Table 3). The LD Score Regression (LDSC) intercept approached one for all phenotypes, indicating limited bias from population stratification. The robustness of discovery GWAS results is illustrated by polygenic score prediction and lead SNP validation (Supplementary Methods 1.3-1.4) in a replication sample (Supplementary Results 2.2-2.3).

### Axon guidance and synaptic functioning genes implicated in visual network-SC GWAS

We continued by examining the possible functional consequences of the SNPs involved in RSN-FC and RSN-SC. SNPs in linkage disequilibrium (LD; *r*^2^ ≥ 0.6) with the Bonferroni-corrected genome-wide significant SNPs from the GWAS which also had suggestive *p*-values (< 1×10^-5^) and a minor allele frequency (MAF) > 0.005 were annotated in FUMA v1.3.7^25^. A detailed overview of the functional annotation of all candidate SNPs is displayed in Supplementary Table 4, whereas the mapped genes that resulted from positional, expression quantitative trait loci (eQTL) and chromatin interaction mapping in FUMA are listed in Supplementary Table 5.

For visual network-SC, an exonic nonsynonymous (ExNS) SNP located in exon 1 of *AC007382.1* (rs711244, *p* = 1.42×10^-12^, CADD = 10.39) was among the candidate SNPs in the locus on chromosome 2. The function of *AC007382.1* is unknown, but it has been associated with amygdala volume previously^26^. Within the loci on chromosome 10 and 7, exonic synonymous SNPs were found in exon 7 and exon 12 of *FAM175B* and *SEMA3A* respectively. The transcript of *FAM175B* is a component of the BRISC enzyme complex that deubiquitinates Lys-63 linked chains in order to control protein function^27^. Experimental studies have suggested that such deubiquitination can regulate synaptic transmission and synaptic plasticity^28^. *SEMA3A* contained multiple intronic SNPs associated with visual network-SC with high CADD scores (11 SNPs with CADD > 12.37), which are usually considered reducing organismal fitness and correlating with molecular functionality and pathogenicity^29^. The product of *SEMA3A* is known as a key regulator of axon outgrowth during the establishment of correct pathways in the developing nervous system^30^.

We additionally mapped 46 visual network-SC candidate SNPs to *METTL10,* because of their established eQTL associations in fetal and adult cerebral cortex tissue as well as through chromatin interaction mapping. *METTL10* encodes a methyltransferase that catalyses the trimethylation of eEF1A at Lys-318 – a key regulator of ribosomal translation^31^. Visual network-SC SNPs were also mapped to the *METTL10-FAM53B* readthrough (*RP11-12J10.3)* and *FAM53B* gene, because of known chromatin interaction in fetal and adult cerebral cortex tissue (Figure 2a). FAM53B is required for Wnt signaling, a pathway important for cell regeneration^32^. Lastly, positional mapping of candidate SNPs within a 10kb window of a gene resulted in the identification of *VIT, STRN, and HEATR5B* genes for visual network-SC (Supplementary Table 5).

**Figure 1.**
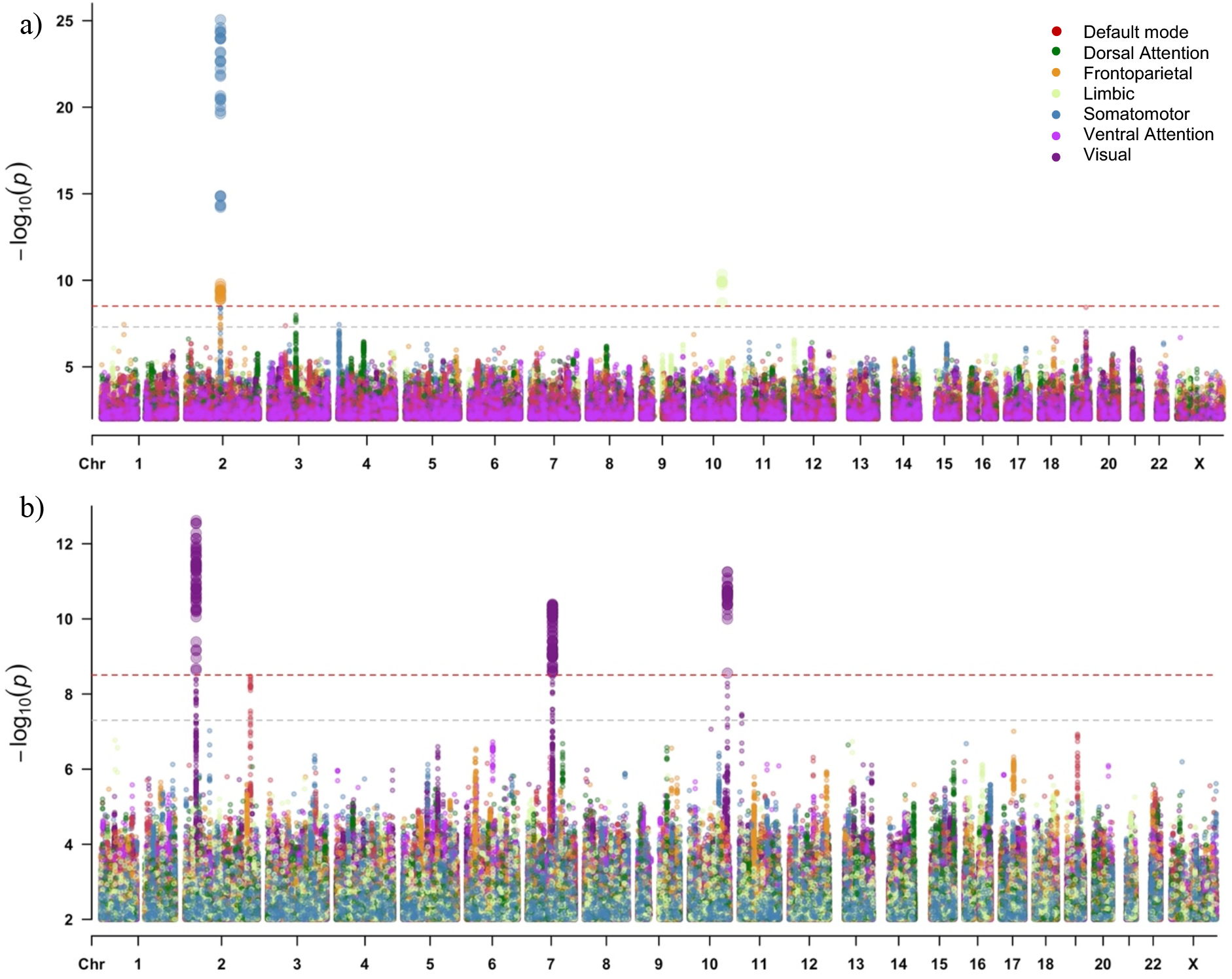
Multitrait Manhattan plots of SNP-based GWAS for a) RSN-FC and b) RSN-SC. The light grey dashed horizontal line indicates traditional genome-wide significance (*p* < 5×10^-8^), whereas the red dashed horizontal line indicates genome-wide significance after additional correction for the number of traits tested (*p* < 3.13×10^-9^). SNPs with *p* > 0.01 are omitted for visualisation purposes. Manhattan plots per RSN are provided as Supplementary Figure 3a (FC) and 4a (SC).

**Figure 2.**
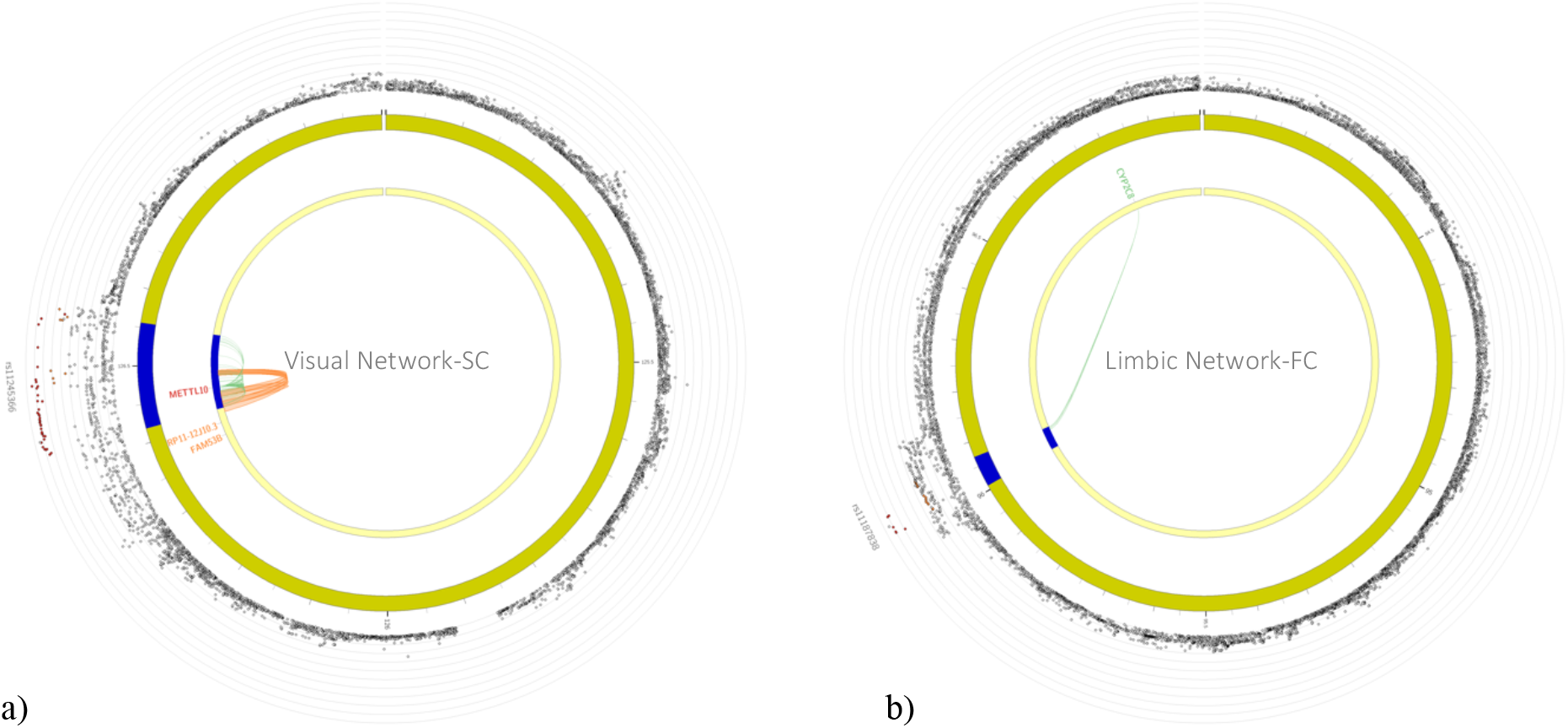
a) Visual network-SC SNPs were mapped to *METTL10, FAM53B* and *METTL10-FAM53B* readthrough (*RP11-12J10.3)* through chromatin interaction mapping (orange). *METTL10* was additionally mapped by 46 SNPs because of their eQTL associations in cerebral cortex tissue. b) FUMA gene mapping, based on established eQTL associations (green) in human temporal cortex, link eight limbic network-FC SNPs on chromosome 10 to *CYP2C8.*

### Annotation of specific and shared loci across RSN-FC

We observed two ExNS SNPs in exon 19 (rs2274224, *p* = 1.771×10^-10^) and 25 (rs2274223, *p* = 1.22×10^-5^) of the *PLCE1* gene to be associated with limbic network-FC. The *PLCE1* gene encodes for the phospholipase C ϵ1, which mediates the production of two second messengers that regulate cell growth, differentiation, and gene expression^33^. The high CADD scores (17.35 and 17.48 respectively) suggest deleteriousness of these two ExNS SNPs. Additionally, four intergenic SNPs within the same locus were located near the *NOC3L* gene.

On chromosome 10, eight SNPs associated with limbic network-FC were eQTLs for the *CYP2C8* gene (Figure 2b). Expression of *CYP2C8* results in an enzyme important for drug metabolism^34^. One of CYP2C8 substrates, the non-selective monoamine oxidase inhibitor phenelzine, is known to target the nervous system and is clinically prescribed as treatment for major depressive disorder^35^. A large body of research has verified the association between major depressive disorder and changes in limbic network functional connectivity, as well as with other RSNs (see Kaiser et al^36^ for a meta-analysis).

The annotation of SNPs in the locus that was shared between frontoparietal and somatomotor network-FC revealed only intergenic candidate SNPs (enrichment = 2.15, *p* = 5.09×10^-9^), which convolutes biological interpretation but is a common observation for complex traits^37^. The nearest genes to the candidate SNPs in this locus were *PAX8* and *IGKV1OR2-108* (respectively 29 and 53 kb distance). *PAX8* encodes a transcription factor that is considered to regulate the expression of genes important for thyroid development^38^ and the production of thyroid hormone^39^. FC within both the somatomotor and frontoparietal network is reduced in individuals with subclinical^40^ and clinical hypothyroidism^41^.

### Default mode network-FC genes associated with Alzheimer’s disease

We next performed gene-based GWAS for the FC and SC within every RSN using MAGMA (Supplementary Table 5). We detected two Bonferroni-corrected genome-wide significant genes additional to the FUMA mapped genes by combining information from neighbouring variants within a single gene in MAGMA (Figure 3, Supplementary Table 6). Visual network-FC was associated with *APOC1 (z* = 5.15, *p* = 1.31×10^-7^), and for default mode network-FC *APOE* was found to be associated (*z* = 5.13, *p* = 1.43×10^-7^). *APOC1* and *APOE* are both located within the 19q13.2 locus and are well-known risk factors for Alzheimer’s disease^42^. Additionally, gene-set analysis results are provided in Supplementary Methods 1.2 and Supplementary Results 2.1.

**Figure 3.**
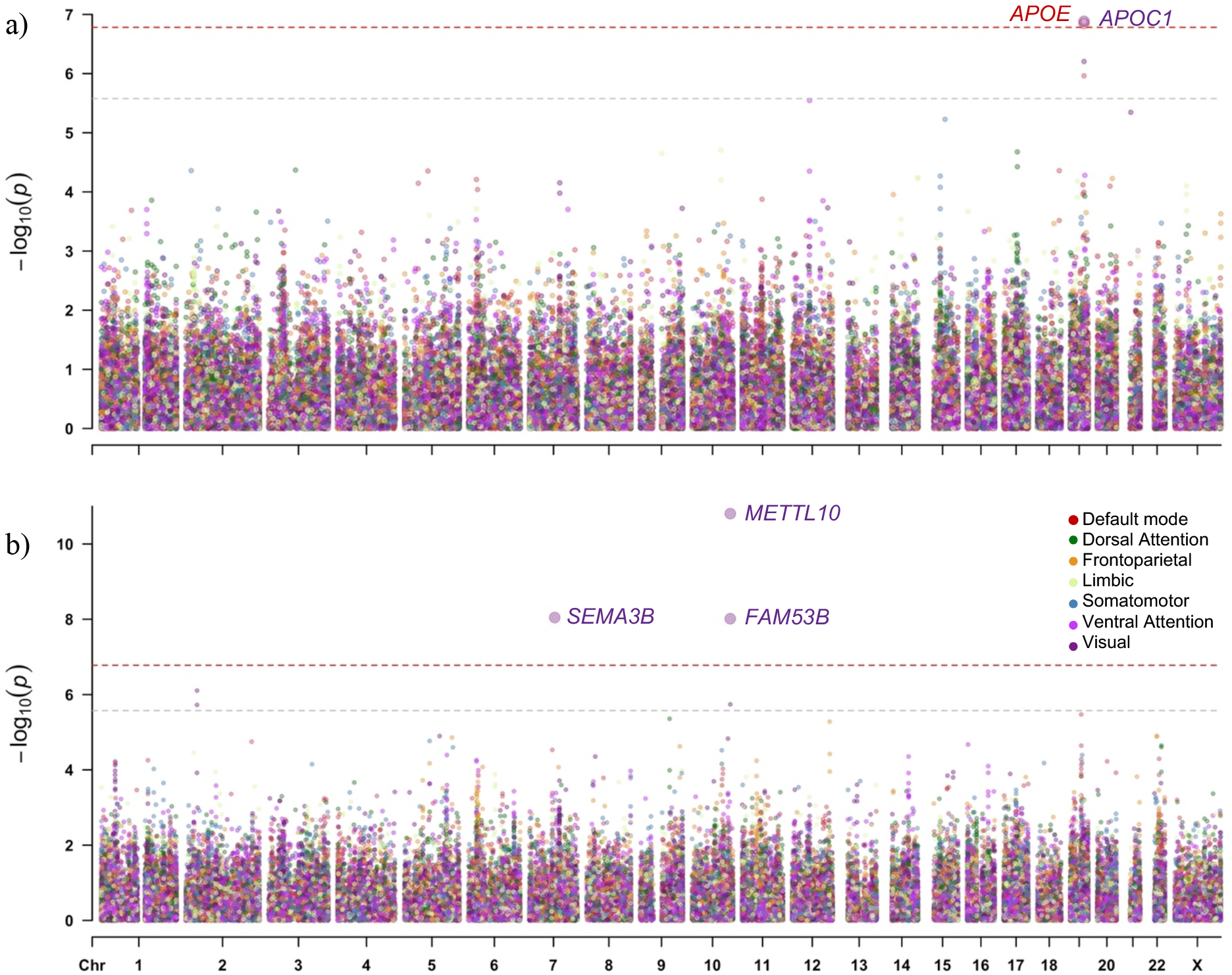
Multitrait Manhattan plots of gene-based GWAS for a) FC and b) SC within RSNs. The light grey dashed horizontal line indicates significance after correcting for the number of genes tested per trait (*p* < 2.65×10^-6^), whereas the red dashed horizontal line indicates significance after an additional correction for the number of traits tested (*p* < 1.66×10^-7^). Manhattan plots per RSN are provided as Supplementary Figure 3b (FC) and 4b (SC).

In order to determine whether there is genetic overlap between Alzheimer’s disease^43^ and default mode network-FC, we performed local genetic correlation (*r*g) analysis using LAVA (see Methods; Supplementary Table 7). For default mode network-FC, we detected two loci on chromosome 12 (BP 64,403,858-66,114,643) and 19 (BP 45,040,933-45,893,307) which showed significant local *r*_g_ at *p* < (0.05/71=) 7.04×10^-4^ with Alzheimer’s disease (Supplementary Figure 5). Given the negligible heritability of global FC in these loci (univariate *p* = 0.27 and *p* = 0.01 respectively, whereas *p* = 1.30×10^-5^ and *p* = 1.62×10^-8^ for default mode network-FC) we conclude that these local genetic associations with Alzheimer’s disease are not driven by total brain connectivity. The locus on chromosome 12 showed a positive *r_g_*(*ρ*) between Alzheimer’s disease and default mode network-FC (BP 64,403,858-66,114,643, *ρ* = 0.69, 95% CI = 0.35 – 1.00, *p* = 3.25×10^-4^). Interestingly, this locus has been identified in a previous GWAS for hippocampal atrophy, a biological marker of Alzheimer’s disease^44^. Negative *r*_g_ between Alzheimer’s disease and FC within DMN was observed in the locus on chromosome 19 (BP 45,040,933-45,893,307, *ρ* = −0.56, 95% CI = −0.82 – −0.38, *p* = 9.23×10^-9^), indicating that lower default mode network-FC was associated with higher genetic risk of Alzheimer’s disease. Note that this larger defined locus showed weak heritability (*p* = 0.014) for visual network-FC despite the significance of *APOC1* in the gene-based GWAS, which would make genetic correlation estimates with Alzheimer’s disease unreliable and uninterpretable^23^. Therefore, Alzheimer’s disease seems to show genetic overlap specifically with default mode network-FC.

### Examining overlap between structure and function per RSN through genetic correlations

As SC strength has been noted to correlate with FC strength on the phenotypic level^16^, we sought to investigate the correlations between FC and SC within each RSN on a genetic level. Genome-wide genetic correlations (*r*_g_) were estimated in LDSC using SNP-based summary statistics (Figure 4). We observed no nominally significant genome-wide *r*_g_’s between SC and FC in any of the RSNs (Supplementary Table 8). Genome-wide *r*_g_ estimates ranged from −0.19 (SE = 0.15, *p* = 0.19) in the dorsal attention network (DAN) and 0.23 (SE = 0.23, *p* = 0.30) in the frontoparietal network (FPN).

**Figure 4.**
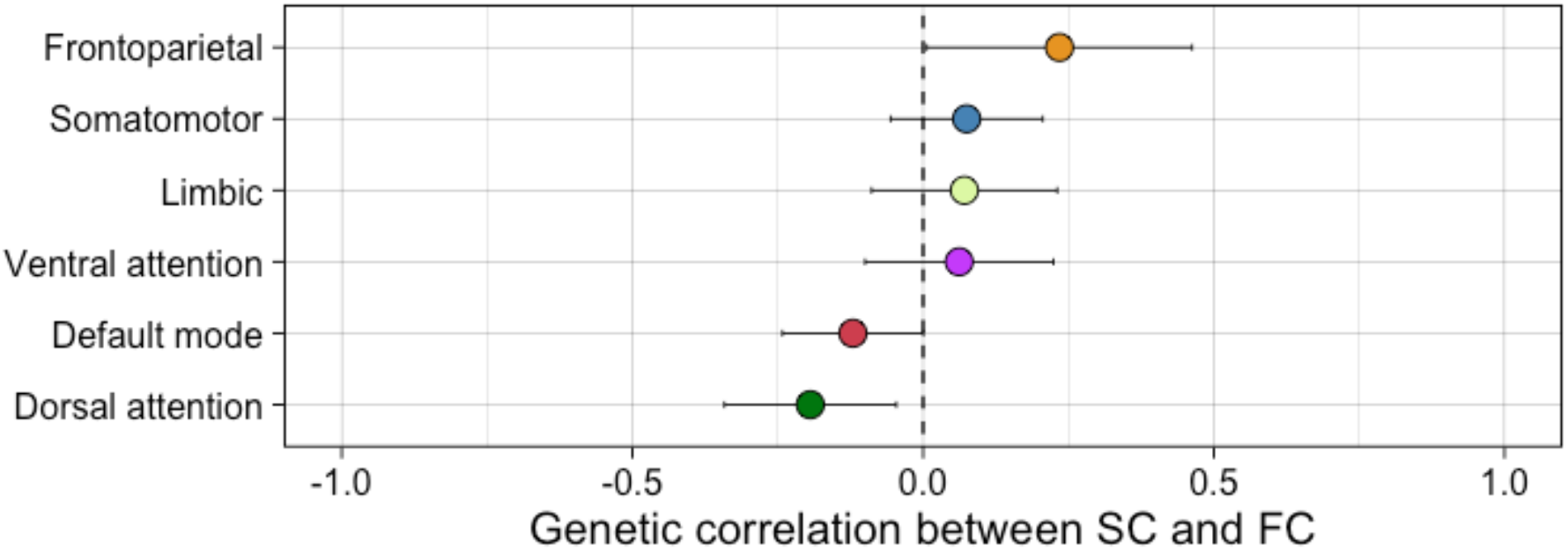
Global *r_g_* (±SE) between FC and SC within the same RSN as performed in LDSC do not show estimates significantly different from zero (Supplementary Table 8). Additional estimation of local *r_g_* did not yield significant overlapping loci between SC and FC within each RSN either (Supplementary Table 9).

Strongly localized or opposing local *r*_g_’s possibly may go undetected, since genome-wide *r*_g_’s are an average of the shared genetic association signal across the genome. We examined whether such relationships between SC and FC within any given RSN exist by performing local *r*_g_ analysis using LAVA^23^, though we did not identify any significant *r*_g_ on a locus level either (Supplementary Table 9).

### Genome-wide and local genetic correlations within the functional and structural domain

We examined the shared genetic signal across RSNs within the same domain by conducting genome-wide *r*_g_ analyses using LDSC (Figure 5; Supplementary Table 8). For functional connectivity, a positive Bonferroni significant genome-wide *r*_g_ was observed between the default mode and ventral attention network (*r*_g_ = 0.52, SE = 0.16, *p* = 1.00×10^-3^). This association was not driven by global FC as neither default mode nor ventral attention network-FC were genetically correlated with global FC (*r*_g_ = 0.19, SE = 0.18, *p* = 0.29; *r*_g_ = 0.26, SE = 0.19, *p* = 0.18 respectively). Note that this positive *r*_g_ does not imply simultaneous functional activation of these two RSNs or their involvement in similar cognitive tasks (which would contradict previous research^45^), but suggests that variants that influence default mode network-FC generally tend to influence ventral attention network-FC in the same direction.

**Figure 5.**
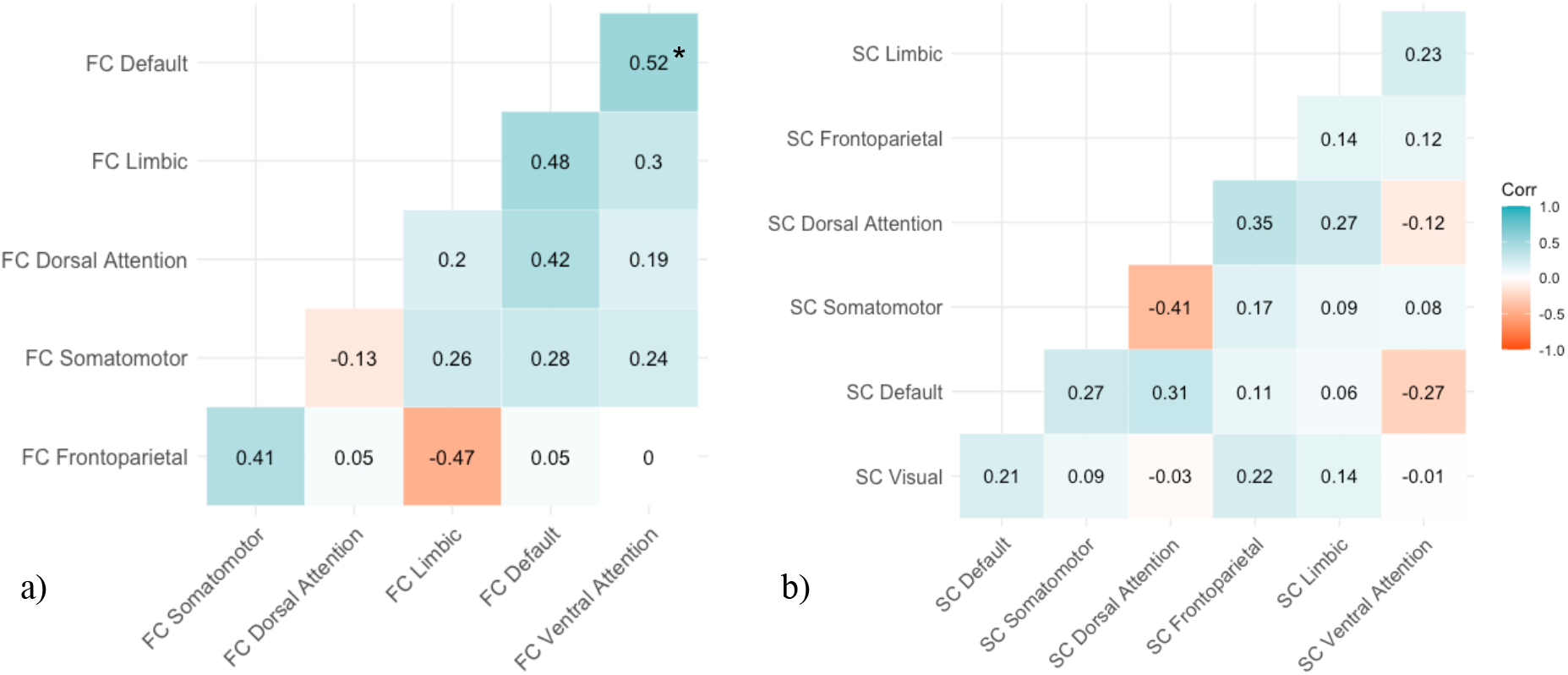
Genome-wide *r_g_* between (a) RSN-FC and (b) RSN-SC. If one of the two RSNs showing significant LDSC *r_g_* showed additional significant *r_g_* with global FC/SC, we instead report the residual *r_g_ (r_g_* between the two RSNs while taking global FC/SC into account in Genomic SEM; see Methods and Figure 6). The significant *r_g_* that survived correction for multiple testing (*p* < 1.19×10^-3^) is indicated with an asterisk (*).

For structural connectivity, we observed multiple significant genome-wide *r*_g_’s (*p* < 1.19×10^-3^) between RSNs, though many of these were also correlated with global SC (Supplementary Table 6). To determine whether the correlations between the structural RSNs could be accounted for by global SC, we used genomic SEM to compute residual *r*_g_ estimates between the structural RSNs while taking global SC into account (see Methods). As none of the residual *r*_g_ estimates remained significant, we conclude that global SC likely accounts for the observed relations between the RSN-SC.

We extended our investigation into shared genetic signal across RSNs beyond the global to the local scale. Eighteen loci showed Bonferroni corrected significant *r*_g_’s when comparing RSNs within the functional domain (Table 1). These were all highly positive (mean *ρ* = 0.84, SD = 0.09) and were not confounded by global FC. When comparing RSNs within the structural domain, local *r_g_* analysis with LAVA revealed only one positively correlated locus between SC within DAN and FPN (15:39238841:40604780, local *r_g_* (*ρ*) = 0.85, *p* = 9.51×10^-7^; Table 1). A complete overview of LAVA local *r_g_* results can be found in Supplementary Table 9.

**Table 1.** Loci with Bonferroni-corrected significant (p< (0.05/774=) 6.46×10^-5^) r_g_ (ρ with lower and upper limit of 95% confidence interval) between RSN-FC or RSN-SC as performed in LAVA. Within these loci, global FC or SC did not show significant univariate h^2^ or r_g_ with either of the two RSNs. See Supplementary Table 9 for all local r_g_ summary statistics. SMN = somatomotor network, VN = visual network, DMN = default mode network, FPN = frontoparietal network, VAN = ventral attention network, DAN = dorsal attention network, LN = limbic network.

| Chr | Start | Stop | RSN 1 | RSN 2 | $\rho$ | 95% CI | | $p$-value |
| --- | --- | --- | --- | --- | --- | --- | --- | --- |
| 1 | 2,215,496 | 2,983,519 | FC SMN | FC VN | 0.77 | 0.47 | 1.00 | $3.39 \times 10^{-5}$ |
| 1 | 18,427,821 | 19,238,924 | FC DMN | FC FPN | 0.72 | 0.45 | 1.00 | $9.48 \times 10^{-6}$ |
| 1 | 211,082,893 | 212,3475,82 | FC VAN | FC SMN | 1.00 | 0.74 | 1.00 | $1.75 \times 10^{-7}$ |
| 2 | 113,930,669 | 115,203,835 | FC FPN | FC SMN | 0.88 | 0.64 | 1.00 | $3.42 \times 10^{-7}$ |
| 2 | 207,726,595 | 208,674,588 | FC FPN | FC VN | 0.97 | 0.72 | 1.00 | $1.02 \times 10^{-6}$ |
| 5 | 4,636,543 | 5,828,694 | FC DMN | FC DAN | 0.73 | 0.47 | 1.00 | $2.72 \times 10^{-5}$ |
| 5 | 68,006,994 | 71,468,651 | FC VAN | FC SMN | 0.79 | 0.53 | 1.00 | $2.05 \times 10^{-5}$ |
| 5 | 75,959,516 | 77,290,255 | FC DMN | FC DAN | 0.91 | 0.65 | 1.00 | $3.42 \times 10^{-6}$ |
| 6 | 10,416,551 | 11,790,671 | FC VAN | FC SMN | 0.83 | 0.55 | 1.00 | $2.49 \times 10^{-5}$ |
| 7 | 50,894,509 | 51,951,647 | FC LN | FC VN | 0.88 | 0.57 | 1.00 | $5.12 \times 10^{-6}$ |
| 8 | 64,215,359 | 66,018,204 | FC DMN | FC VN | 0.86 | 0.59 | 1.00 | $1.09 \times 10^{-5}$ |
| 9 | 93,441,051 | 94,175,374 | FC FPN | FC SMN | 0.90 | 0.61 | 1.00 | $1.73 \times 10^{-5}$ |
| 9 | 93,441,051 | 94,175,374 | FC FPN | FC VN | 0.87 | 0.62 | 1.00 | $4.58 \times 10^{-6}$ |
| 10 | 89,971,629 | 91,021,321 | FC VAN | FC VN | 0.96 | 0.67 | 1.00 | $1.23 \times 10^{-6}$ |
| 15 | 39,238,841 | 40,604,780 | SC DAN | SC FPN | 0.85 | 0.53 | 1.00 | $9.51 \times 10^{-7}$ |
| 17 | 13,648,447 | 14,508,610 | FC DMN | FC LN | 0.89 | 0.69 | 1.00 | $3.50 \times 10^{-9}$ |
| 18 | 2,839,843 | 3,722,828 | FC DMN | FC DAN | 0.70 | 0.45 | 1.00 | $2.66 \times 10^{-5}$ |
| 19 | 17,045,964 | 17,750,518 | FC LN | FC DAN | 0.73 | 0.47 | 1.00 | $2.34 \times 10^{-6}$ |
| 19 | 17,045,964 | 17,750,518 | FC DMN | FC DAN | 0.79 | 0.53 | 1.00 | $1.43 \times 10^{-5}$ |

## Discussion

Mapping the genetic components of resting state networks (RSNs) may provide insight into the aetiology of brain function and brain disorders. RSNs are typically defined using functional connectivity (FC), and structural connectivity (SC) correlates to FC in varying degrees across RSNs^17^. The genetic component of RSN-SC has been less studied and as one of the fundamental goals in neuroscience is to understand the relationship between structure and function within the brain, the aim of this study was to gain more insight into the genetic underpinnings of structural and functional connectivity (SC; FC) within a framework that respects the brain’s hierarchical functional architecture. With the use of GWAS and in silico annotation we identify the first genes for visual network-SC, that are involved in axon guidance and synaptic functioning. We further observe that genetic variation in RSN-FC (e.g. limbic network-FC and default mode network-FC) impacts biological processes related to brain disorders (major depressive disorder and Alzheimer’s disease respectively) that have previously been associated with FC alterations in those same RSN. The genetic component of RSNs overlaps mostly within the functional domain, whereas less overlap is observed within the structural domain and between the functional and structural domains.

For FC within RSNs (RSN-FC), we detect biologically interpretable results that are specific to default mode and limbic network-FC. For default mode network-FC, we observe *APOE* as a genome-wide significant gene. The default mode network is hypothesized to relate to Alzheimer’s disease through the role of default model network-FC in memory consolidation^46^ and through the spreading of cortical atrophy over time, which follows the pattern of default mode network regions^47^. Here, we complement earlier phenotypic observations that link Alzheimer’s disease to default mode network-FC^48^ by now also showing genetic correlations in two loci between Alzheimer’s disease and default mode network-FC. Functional follow up would be necessary to investigate how the variants and genes in these loci affect default mode network-FC. The limbic network is commonly known for its involvement in emotion regulation, episodic memory, and action–outcome learning^49^ and has been associated with mood disorders, such as major depression disorder and bipolar depression^50^. The genes *PLCE1, NOC3L* and *CYP2C8* were related to limbic network-FC, all of which have been noted to have a relationship with major depressive disorder^35,51,52^. A previous study investigating the role of *PLCE1* in major depressive disorder patients has demonstrated an association with antidepressant remission in female patients, together with other genes within the calcium/calmodulin-dependent protein kinase (CaMK) pathway^51^. *NOC3L* eQTLs in the cerebellum and nucleus accumbens have previously been demonstrated to associate with depression severity and antidepressant response^52^, and one of the substrates of CYP2C8 is clinically prescribed as treatment for major depressive disorder (phenelzine)^35^. These results seem to suggest that major depressive disorder and antidepressant response involve processes that are impacted by genetic variation in limbic network-FC.

In addition to RSN-FC specific effects, we find evidence of shared genetic signal in FC across different RSNs using several approaches. Specifically, we observe a genetically correlated and common genome-wide significant locus for both somatomotor and frontoparietal network-FC near *PAX8. PAX8* regulates multiple genes involved in the production of thyroid hormone^39^, an interesting result considering that both somatomotor and frontoparietal network-FC have been linked to (subclinical) hypothyroidism^40,41^. Additionally, we detect genetically correlating loci between all RSN-FC and a genome-wide genetic correlation between ventral attention and default mode network-FC. The ventral attention network supports salience processing^53^, whereas the default mode network includes areas widespread over the brain and supports emotional processing, self-referential mental activity, and recollection of prior experiences^54^. Increased FC within these two RSNs has been associated with bulimia nervosa^55^ and contributes to episodic memory retrieval^53^. Altogether, the shared genetic underpinnings of different RSN-FC that we present here could give a possible explanation how multiple disorders are associated with more than one RSN.

We report considerable heritability estimates for RSN-SC (ranging from 10.00% to 15.40%) and identify nine genes that suggest a role for synaptic transmission in the genetics of visual network-SC. For example, *STRN* encodes for a calmodulin-binding protein that is mostly found in dendritic spines playing a role in Ca2+-signaling^56^, the transcript of *FAM175B* is a component of a deubiquitylation enzyme complex that has been suggested play a role in synaptic transmission and synaptic plasticity^28^, and *SEMA3A* is known as an axonal guidance gene during development^30^. The SEMA3A protein has been shown to be upregulated in schizophrenia patients and is suggested to contribute to the developmentally induced impairment of synaptic connectivity in the disorder^57^. Visual network functional hyperconnectivity has been observed in schizophrenia^58,59^ and related to visual hallucinations^59^, but future studies should investigate the equivalent SC component in more detail given our findings.

When investigating the genetic relationship between SC and FC within each RSN, we find no significant genome-wide or local genetic correlations. Since the estimation of genetic correlations is dependent on sample size and the heritability estimates of both traits^60^, studies with increased power are needed to examine the robustness of these results. Future studies could additionally incorporate recent insights that indirect structural connections supporting direct functionally connected regions complicate simple structure to function mapping^61^. Our study focussed on direct structural connections within RSNs. The possibility that the genetics of RSN-FC overlap with that of indirect pathways that structurally connect brain regions within RSNs via a route beyond the borders of that RSN could therefore be subject to future research.

Several limitations must be considered while interpreting our results. It is known that rsfMRI measures can be noisy and subject to motion distortion, which raises the possibility of differences in measurement error between RSN-FC and RSN-SC. However, given our stringent pre-processing and quality control to enable noise minimization and additional use of rsfMRI-specific covariates in GWAS, we were able to find heritability estimates for RSN-FC that are concordant with previous studies^13^. Second, even though UK Biobank provides genetic and uniform MRI data at unprecedented sample sizes, it is evident that even larger sample sizes are needed for discovering the often small genetic effects of polygenic traits^62^. The null results observed for some RSN-FC/SC GWAS, partitioned heritability and gene-set analyses might be explained by the multiple comparison correction for the number of phenotypes analysed, in conjunction with insufficient statistical power. Third, some other sample characteristics, such as the European ancestry, age-class and socioeconomic status of subjects, may limit the generalizability of our findings. While we corrected for age and Townsend deprivation index (a proxy of socio-economic status) in our GWAS to reduce this bias, larger and more diverse imaging-genetics datasets are undoubtedly needed.

This study examines the specificity and overlap in genetic architecture of RSNs – structurally and functionally. We observe several genetic effects that seem to be specific to certain RSNs and highlight relevant biological processes for brain connectivity and related brain disorders. The complexity of structure-function coupling within RSNs is illustrated by the observation that, despite genetic overlap of RSNs within the functional domain, genetic overlap is less apparent within the structural domain and between the functional and structural domains. Altogether, this study advances the understanding of the complex functional organisation of the brain and its structural underpinnings from a genetics viewpoint.

## Methods

A flowchart that describes all Methods used in this manuscript is displayed in Supplementary Figure 1.

### Sample

The UK Biobank (UKB) is a resource with genomic and imaging data of volunteer participants^63^. The National Research Ethics Service Committee North West–Haydock ethically approved this initiative (reference 11/NW/0382) and data were accessed under application #16406. Combined SNP-genotypes and neuroimaging data of N = 40,682 participants have been available since January 2020. From all new subjects ID’s in the latest neuroimaging release (January 2020), we randomly assigned 5,000 subjects to a holdout set for validation. Subsetting the total sample to subjects with all neuroimaging data necessary to construct our phenotypes as described below, resulted in N_FC_ = 37,017 and N_SC_ = 36,645. We only included subjects for which the projected ancestry principal component score was closest to and < 6 SD from the average principal component score of the European 1000 Genomes sample based on Mahalanobis distance. This procedure has been described in previous publications by our group^64^ and the number of non-European exclusions are displayed in Supplementary Table 10. Other exclusion criteria were withdrawn consent, UKB-provided relatedness, discordant sex or sex aneuploidy (Supplementary Table 10). Further quality control on genomic and neuroimaging data is described below and resulted in the sample sizes and sample characteristics as displayed in Supplementary Table 11.

### Genotype data

The genotype data used in this study were obtained from the UK BiobankTM Axiom and the UK BiLEVE Axiom arrays. These Affymetrix arrays cover 812,428 unique genetic markers and overlap 95% in SNP content. This number of SNPs was increased to 92,693,895 by imputation carried out by UKB. Variants were imputed using the Haplotype Reference Consortium and the UK10K haplotype panel as reference. We applied our in-house quality control pipeline in addition to quality control performed by UKB. This procedure excluded SNPs with low imputation scores (INFO<0.9), low minor allele frequency (MAF<0.005) or high missingness (>0.05), multiallelic SNPs, indels, and SNPs without unique rs-identifiers. A total of 9,380,668 SNPs passed quality control and were converted to hard call SNPs using a certainty threshold of 0.9 for further analyses.

### Neuroimaging data

#### Pre-processing & connectome reconstruction

The UKB scanning protocol and processing pipeline is described in the UKB Brain Imaging Documentation^65^. For this study, we made use of the available resting-state functional brain images (rsfMRI) and multiband diffusion brain images (DWI) together with T1 surface model files and structural segmentation from FreeSurfer^66^. These three types of data were used as input for the structural and functional pipeline of CATO (Connectivity Analysis TOolbox)^67^. Prior to this, UKB performed pre-processing on DWI and rsfMRI data as described in the UKB Brain Imaging Documentation^65^.

In CATO’s structural pipeline, additional pre-processing of DWI files was performed in FSL^68^ by computing a DWI reference image based on the corrected diffusion-unweighted (b0) volumes, computing the registration matrix between DWI reference image and the anatomical T1 image, and registering the Freesurfer segmentation to the DWI reference image. The surface was parcellated based on the Cammoun sub-parcellations of the Desikan-Killiany atlas including 250 cortical regions^69^. We reconstructed the diffusion signal with diffusion tensor imaging (DTI), a deterministic method that is robust and relatively simple compared with more advanced diffusion reconstruction methods^67^. In CATO, the Fiber Assignment by Continuous Tracking (FACT) algorithm^70^ is used to reconstruct fibers and fractional anisotropy (FA) was used as weights of reconstructed fibers. FA is a robust measure of white matter integrity and has been found to be sensitive to changes in connectivity^18^ and correlates with axon density, size and myelination^71^. The structural connectivity matrix was built out of all fiber segments that connected two regions in the atlas. Additional filters were applied, namely a minimal FA of 0.1, minimal length of 30 mm and having 2 or more number of streamlines.

The functional pipeline in CATO consisted of similar steps. First, we computed a rsfMRI reference image by averaging all rsfMRI frames in FSL and subsequently registered this reference image and the T1 image in FreeSurfer. Second, we parcellated the surface based on the same atlas as in the structural pipeline (to enable structure-function comparison in downstream analyses) and we registered the T1 parcellation to the rsfMRI image. Third, motion metrics were estimated, and time-series were corrected for covariates (linear trends and first order drifts of motion parameters and the mean signal intensity of voxels in white matter and cerebrospinal fluid and of all voxels in the brain) by regression. Fourth, time-series were passed through band-pass filtering (frequencies 0.01 to 0.1) and scrubbing (max FD = 0.25, max DVARS = 1.5, min violations = 2, backward neighbours = 1, forward neighbours = 0). Fifth, the functional connectivity matrix was computed by the Pearson’s correlation coefficient of the average signal intensity of every pair of brain regions across the frames that survived filtering.

#### Quality control

The UKB scanning and pre-processing protocol includes filters for outliers based on manual QC and an advanced classifier described elsewhere^72^. We excluded a small number of subjects that UKB identified as outliers and placed in an “unusable” folder. The UKB main documentation^65^ suggests a second set of UKB data fields that can be used as outlier criteria. Outlier subjects are defined as subjects that score for any of the values > 3 interquartile ranges above the upper quartile or below the lower quartile. Outlier criteria included measures that describe the discrepancy between the T1-weighted, rsfMRI and DWI images and the population average template after LINEAR and NON-LINEAR alignment, the amount of nonlinear warping necessary to map a subject to the standard template, the signal to noise ratio in rsfMRI, the mean rfMRI head motion averaged across space and time points and the total number of outlier slices in DWI volumes. We extended this recommended list with connectome specific measures, including the average prevalence of all connections present and absent in the reconstructed brain network of a subject (low average prevalence scores indicate the presence of odd connections and high values indicate the absence of common connections), the sum of number-of-streamlines and average FA of all connections in the reconstructed brain network of a subject. The number of exclusions can be viewed in Supplementary Table 10.

#### Phenotype reconstruction

In this study, the phenotypes of interest were the functional and structural connectivity (FC;SC) within seven resting-state networks (RSNs) that previously have been identified^2^ and are commonly used in (clinical) neuroimaging studies: the default mode network, ventral attention network, dorsal attention network, visual network, limbic network, somatomotor network and frontoparietal network. Each of the 250 cortical regions of the reconstructed structural and functional connectomes were assigned the ratio to what extent they belonged to each of these seven RSNs, using a mask created and validated elsewhere (see Supplementary Information of Wei *et al*^24^). Each connection was then weighted by multiplying the ratios of the two regions involved in the particular RSN. FC and SC within the RSNs were respectively calculated as the mean correlation and mean fractional anisotropy of the connections within the RSN. We also computed two global FC and SC phenotypes as the mean correlation and mean fractional anisotropy of all available connections, to be able to correct for connectivity that is non-specific to RSNs in downstream analyses.

### Statistical analyses

#### SNP-based GWAS

To identify common genetic variants involved in FC within each of the seven RSN, we performed seven SNP-based GWAS in PLINK2^73^. Also, for the SC within each of the seven RSN, a SNP-based GWAS was performed. It is common practice to include a global FC or SC estimate as covariate in GWAS to capture associations that are driven by the level of connectivity within an RSN irrespective of the level of connectivity throughout the whole brain. It has become apparent that this risks the introduction of collider bias (inducing false-positives)^74^. Here we build upon recent developments in statistical genetics that have provided multiple methods that allow for post-GWAS analyses conditional on global connectivity. Therefore, we used the global FC and global SC phenotypes to run two additional SNP-based GWAS, for which the summary statistics were used for conditional downstream analyses. The total amount of GWAS was therefore sixteen. In order to correct for population stratification during GWAS, a principal component analysis was performed in FlashPCA2^75^ using only independent (*r*^2^ < 0.1), common (MAF > 0.01) and genotyped SNPs or SNPs with very high imputation quality (INFO=1). The first 30 principal components were used as covariates in all GWAS, together with sex, age, genotype array, Townsend deprivation index (a proxy of socio-economic status), general neuroimaging confounders as well as FC/SC specific covariates (recommended by Alfaro-Almagro and colleagues^76^). The general set included handedness, scanning site, the use of T2 FLAIR in Freesurfer processing, intensity scaling of T1, intensity scaling of T2 FLAIR, scanner lateral (X), transverse (Y) and longitudinal (Z) brain position, and Z-coordinate of the coil within the scanner. FC-specific and SC-specific covariates were respectively intensity scaling and echo time of rsfMRI, and intensity scaling of DWI. For reasons of collinearity, we ran principal component analysis on all covariates (excluding the population stratification principal components) and retained those principal components that explained > 99% of variance. Rare variants (MAF < 0.005) and SNPs with high missingness (>5%) were excluded from GWAS and male X variants were counted as 0/1. The genome-wide significance threshold was α = (0.05/1,000,000/16 =) 3.13×10^-9^ according to the Bonferroni correction for multiple testing.

#### SNP-based heritability

SNP-based (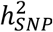; or narrow-sense) heritability represents the proportion of phenotypic variance that can be explained by common additive variation. In contrast, broad-sense heritability captures the total genetic contribution to the phenotype and is often based on family studies^77^. We applied Linkage Disequilibrium Score regression (LDSC) on the SNP-based GWAS summary statistics of all sixteen phenotypes to estimate 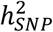 using precomputed LD scores from 1000 Genomes EUR, as provided by the LDSC developers.

#### Functional annotation

FUMA is a web-based platform that can be used to functionally map and annotate SNPs that appear significant during GWAS. We uploaded summary statistics to FUMA if GWAS identified at least one genome-wide significant SNP. Candidate SNPs were defined as all SNPs in LD *r*^2^>0.6 with an independent genome-wide significant SNPs (*r*^2^<0.6). Annotation was subsequently performed using ANNOVAR^78^, RegulomeDB^79^ score and ChromHMM^80^. Lead SNPs were defined as independent SNPs *r*^2^<0.1. Genomic loci were constructed by taking all independent significant SNPs *r*^2^ < 0.1 with LD blocks within 250 kb distance and independent significant SNPs *r*^2^ ≥ 0.1. Within every locus, SNPs were mapped to genes using three methods: positional mapping, eQTLs mapping or chromatin interaction mapping. SNPs were positionally mapped to genes if their physical distance was <10 kb. Mapping based on eQTLs relied on known associations between SNPs and the gene-expression of genes within a 1Mb window, from BRAINEAC^81^ (frontal, occipital, temporal, cerebral cortex), GTEx v8^82^ cerebral cortex and xQTLServer^83^ dorsolateral prefrontal cortex. Chromatin interaction mapping was based on established 3D DNA-DNA interactions between SNP and gene regions from Hi-C databases in cortex tissue (PsychENCODE^84^, Giusti-Rodriguez *et al*^85^, and GSE87112^86^). To restrict chromatin interaction mapping to plausible biological interactions, we only included interactions where one region overlapped with an enhancer (as predicted by the Roadmap Epigenomics project^87^ in cortex tissue) and the other region overlapped with a promoter (250 bp upstream to 500 bp downstream of the transcription start site as well as predicted by the Roadmap Epigenomics project in cortex tissue). A FDR threshold of 1×10^-5^ was used, as recommended in previous literature^86^.

#### Gene-based GWAS

Performing GWAS on the level of genes has been suggested to be more powerful than GWAS on the level of SNPs^88^. Therefore, the sixteen SNP-based GWAS summary statistics were used to perform sixteen gene-based GWAS in MAGMA (Multi-marker Analysis of GenoMic Annotation) v1.08^88^. A mean SNP-wise model was applied (with the UKB European population serving as an ancestry reference group) to test the joint association of all SNPs within 18,850 genes with RSN-FC/RSN-SC. The genome-wide significance threshold was adjusted for multiple testing to α = (0.05/18,850)/16 = 1.66 ×10^-7^.

#### Genome-wide genetic correlations

To assess the overlap in genetic architecture between FC/SC within RSNs while taking the influence of global FC/SC into account, we designed a genetic correlation (*r*_g_) analysis pipeline. This pipeline consisted of three steps. 1) In the first step, genome-wide *r*_g_ between 42 combinations of RSNs were estimated using LDSC (α= (0.05/42=) 1.19×10^-3^). The summary statistics of SNP-based GWAS were used as input for LDSC. We excluded FC-VN, because both the lambda (<1.02) and ratio (>0.20) values were out of bound for LDSC. 2) For all RSNs included in a significant bivariate *r_g_,* additional *r*_g_ with global FC/SC were calculated in LDSC. 3) If one or both RSNs from the significant bivariate *r*_g_ showed additional significant *r*_g_ with global FC/SC, we recalculated of the genome-wide *r*_g_ between the two RSNs with global FC/SC taken into account. Since such residual genome-wide *r*_g_ analyses are not implemented in LDSC, we applied Genomic Structural Equation Modelling (genomic SEM)^89^. Genomic SEM is a method that enables to model the multivariate genetic architecture and covariance structure of complex traits using GWAS summary statistics and allows for sample overlap. We modelled residual covariance between RSN as the covariance between the residual variance of the two RSNs involved after taking the global factor into account (Figure 6). A confirmatory factor analysis was then ran using Diagonally Weighted Least Square estimation.

**Figure 6.**
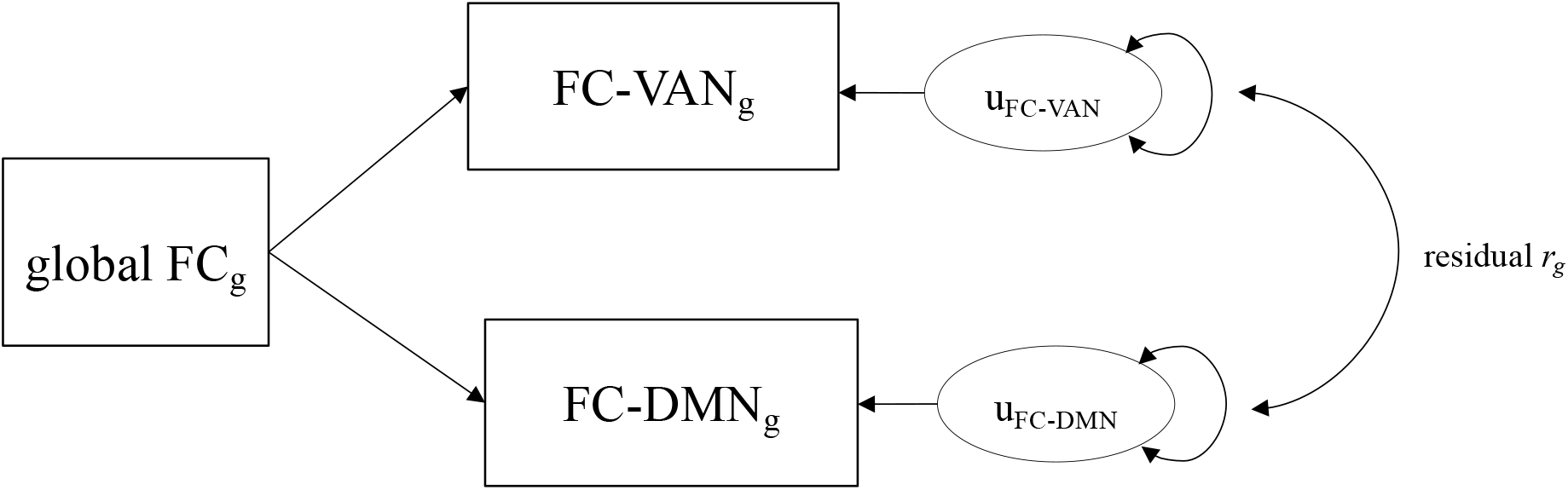
Path diagram of genomic SEM model. The summary statistics of two RSNs that have shown to significantly correlate with global connectivity will be used as input together with summary statistics of the global connectivity GWAS. In this way, *r_g_* between the two RSNs can be estimated while taking global connectivity into account.

#### Local genetic correlations

The genome-wide *r*_g_’s described above are an average correlation of genetic effects across the genome, implicating that contrasting local *r*_g_’s are possibly cancelling each other out. Running *r*_g_ analysis on a locus level has the potential to uncover loci that show genetic similarity between traits. For this purpose, we adopted a three-step local *r*_g_ analysis pipeline similar to the genome-wide *r*_g_ analysis approach described above. All three steps were performed in LAVA^23^, a local *r*_g_ analysis tool, using SNP-based GWAS summary statistics as input. We followed the suggested sample overlap procedure (as described on https://github.com/josefin-werme/LAVA) to enable LAVA to model shared variance due to sample overlap as residual covariance and consequently remove upward bias in local *r*_g_ estimates^23^. Since our GWASs included European samples, the 1,000 Genomes Phase 3 European data served as genotype reference and formed the basis of the locus definition file. For every locus, the first step of our pipeline consisted of estimating local bivariate *r*_g_ between 49 combinations of RSNs. However, RSNs that were devoid of heritable signal (*p* > 1×10^-4^) in the locus were excluded from local bivariate *r*_g_ analysis to ensure interpretability and reliability. A total of 774 bivariate tests were performed across 337 loci, leading to an adjusted significance threshold of α= (0.05/774=) 6.46×10^-5^. In the second step, RSNs that showed significant local *r*_g_ were additionally tested for *r*_g_ in that locus with global FC/SC. Note that if this was not possible, because global FC/SC showed no significant heritability in that locus, the local bivariate *r*_g_ between RSNs could not be biased by global FC/SC. If one or both RSNs did show additional significant *r*_g_ with global FC/SC, we ran a partial local *r*_g_ between the RSNs conditioned on the SC-global and/or FC-global phenotype in step three. If the partial local *r*_g_ between the RSNs no longer remained significant, we concluded that the initial *r*_g_ was driven by global FC/SC and did not reflect genetic overlap specific for these RSNs.

## Acknowledgements

D.P. was funded by The Netherlands Organization for Scientific Research (NWO VICI 453-14-005), NWO Gravitation: BRAINSCAPES: A Roadmap from Neurogenetics to Neurobiology (Grant No. 024.004.012), and a European Research Council advanced grant (Grant No, ERC-2018-AdG GWAS2FUNC 834057). The work of S.L. was supported by ZonMw Open Competition, project REMOVE 09120011910032. C.A.d.L. is funded by Hoffman-La Roche. The work of M.H. was supported by a VIDI (452-16-015) grant from the Netherlands Organization for Scientific Research (NWO) and an ERC Consolidator of the European Research Council (101001062). J.E.S. was supported by a VENI (201G-064) grant from the NOW. The research has been conducted using the UK Biobank Resource (application no. 16406). Analyses were carried out on the Genetic Cluster Computer hosted by the Dutch National computing and Networking Services SURFsara.

## Author contributions

E.P.T., M.P.v.d.H. and D.P. conceived of the study, and J.W., M.N. and C.A.d.L. contributed additionally to its design. J.E.S. performed preprocessing of genetic data, while E.P.T. and S.C.d.L. performed preprocessing of neuroimaging data together. E.P.T. performed the analyses with contributions from J.W. and Y.W. E.P.T., J.W., S.C.d.L., M.N., C.A.d.L., M.P.v.d.H. and D.P. contributed to the interpretation of the data. E.P.T. drafted the work. E.P.T., J.W., M.N., D.P. and M.P.v.d.H. substantively revised it and S.C.d.L., J.E.S., Y.W. and C.A.d.L. revised it.

## Competing interests

The authors report no competing interests.

## Data availability

Genome-wide summary statistics will be made publicly available via https://ctg.cncr.nl/software/summary_statistics/ upon publication.

## Notes

### Competing Interest Statement

The authors have declared no competing interest.

