## Supplementary Information for "Specificity and overlap in the genetic architectures of functional and structural connectivity within cerebral resting-state networks"

*1. Supplementary Methods*
- *1.1 Partitioned SNP-based heritability……………………………………………………….1*
- *1.2 Gene-set analysis……………………………………………………………………….....2*
- *1.3 Polygenic score…………………………………………………………………………..2*
- *1.4 Internal validation of lead SNPs………………………………………………………...2*

*2. Supplementary Results*
- *2.1 Conditional gene-set analysis for biological pathways……………………………......3*
- *2.2 Polygenic score prediction…………………………………………………………......4*
- *2.3 Internal validation of lead SNPs………………………………………………………....4*

*3. Supplementary Figures*
- *3.1 Figure S1………………………………………………………………………………......5*
- *3.2 Figure S2………………………………………………………………………………......6*
- *3.3 Figure S3………………………………………………………………………………......7*
- *3.4 Figure S4………………………………………………………………………………......8*
- *3.5 Figure S5………………………………………………………………………………......9*

*4 References*

### 1. Supplementary Methods

#### *1.1 Partitioned SNP-based heritability*

For the phenotypes with enough polygenic signal to run LDSC (lambda > 1.02), we investigated if certain functional categories in the human genome were enriched for $h^2_{SNP}$. The ratio of the proportion of $h^2_{SNP}$ in a certain category to the proportion of SNPs in the category equalled the enrichment value. We corrected the level of significance for multiple testing to $\alpha = (0.05/28)/11 = 1.62\times10^{-4}$.

*1.2 Gene-set analysis*

We set out to prioritize the associations from gene-based GWAS and gain more insight into the biological pathways associated to RSN-FC/RSN-SC. In order to identify gene-sets specific for RSN-FC/RSN-SC, we ran conditional gene-set analyses in MAGMA conditioning on the global FC or SC respectively. Pathways were represented as gene-sets from Gene Ontology (GO) molecular functions, cellular components and biological processes, and curated gene-sets from MsigDB v7.0 (sets C2 and C5)[1]. Protein-coding genes served as background genes. The threshold for gene-sets reaching significance was corrected for multiple testing to $\alpha = (0.05/7246)/14 = 4.93\times10^{-7}$.

*1.3 Polygenic score*

The variance explained in RSN-FC/RSN-SC by our GWAS findings was investigated to test the robustness of our findings using polygenic score (PGS) estimation in PRSice-2[2]. We applied a two-phase approach to obtain *p*-values unaffected by overfitting and therefore split our holdout sample in a target (N = 1,818) and validation set (N = 1,824). In phase 1, SNP-based summary statistics (MAF > 0.1, chromosome X excluded) from the discovery sample together with genotype data of the target set were used to find the optimal *p*-value threshold. PRSice-2 uses high-resolution thresholding and clumping of genotype data, and we included the same covariates as during discovery GWAS. In phase 2, the model with the best fit from phase 1 was applied on the validation set to obtain the variance explained ($R^2$).

*1.4 Replication of lead SNPs*

In the design of this study, a hold-out sample of $N_{FC} = 3{,}408$ and $N_{SC} = 3{,}412$ was reserved for PRS analysis (described above). We applied an earlier described method (Okbay et al[3], Supplementary Information 1.8) to internally validate our discovery lead SNPs. This formula describes the probability of a discovery lead SNP *i* being significant in a replication sample as

$$P(sig_i) = \Phi\left(-\frac{|\beta_i|}{\sigma_{rep,i}} + \Phi^{-1}\left(\frac{\alpha}{2}\right)\right) + \left[1 - \Phi\left(-\frac{|\beta_i|}{\sigma_{rep,i}} - \Phi^{-1}\left(\frac{\alpha}{2}\right)\right)\right]$$

with α representing an alpha level of 0.05, $\Phi$ the cumulative normal distribution function, $\Phi^{-1}$ the inverse normal distribution function, $\sigma_{rep,i}$ the standard error of SNP $i$ in the replication GWAS and $\beta_i$ the winner's curse adjusted association estimate of SNP $i$. Winner's curse is the occurrence of overestimated effect sizes that are induced by significance thresholding[4]. We applied winner's curse correction using the mean of the normalized conditional likelihood[5] in the *winnerscurse* R package. The number of SNPs that is expected to show significance was then summed across all six lead SNPs by $\sum_i P(sig_i)$. Given the small effect sizes of GWAS SNPs, a larger sample size is often needed to replicate findings. Since the standard error of a SNP is dependent on sample size, we calculated $P(sig_i)$ for a range of sample sizes by $\frac{SD_{rep,i}}{\sqrt{N}}$ and plotted $P(sig_i)$ across this range to describe the power to replicate these lead SNPs.

### 2. Supplementary Results

#### *2.1 Conditional gene-set analysis on biological pathways*

We looked for convergence of the genetic signal for RSN-FC/RSN-SC onto 7,252 MSigDB[1] pathways using MAGMA gene-set analysis, a useful method for further functional interpretation[6]. We conditioned our analyses on the gene-based GWAS summary statistics for global FC/SC in an effort to capture RSN-specific pathways. Five pathways showed an association with four RSNs after Bonferroni correction for the number of pathways tested per trait (Supplementary Table 12 displays the associations of all pathways tested for all traits). These included blood vessel morphogenesis (GO, $p = 3.30\times10^{-6}$) and vasculature development (GO, $p = 4.94\times10^{-6}$) pathways for SC within LN, the Parkinson's Disease pathway (KEGG, $p = 2.10\times10^{-6}$) for SC within SMN, the pathway for positive regulation of mesenchymal cell proliferation (GO, $p = 2.64\times10^{-6}$) in FC within DMN, and the pathway for negative regulation of histone methylation (GO, $p = 5.23\times10^{-6}$) for the FC within DAN. These five pathways did however not survive further Bonferroni correction for the number of traits tested ($\alpha = (0.05/7{,}246)/14 = 4.93\times10^{-7}$). Therefore, it cannot be concluded that these biological processes are involved in the genetics of FC and SC within RSNs.

*2.2 PGS prediction*

The variance that could be explained in RSN-FC/RSN-SC by polygenic scores based our GWAS associations was considered. In PRSice-2, the summary statistics of the discovery GWAS were used to find the best $p$-value threshold for polygenic scores in the target set. The application of this optimal prediction model in our validation set explained on average 0.28% and 0.35% of the variance across RSN-FC and RSN-SC respectively (Supplementary Table 14). Note that this $R^2$ value is on average 2.75% (SC) to 6.89% (FC) of the $h^2_{SNP}$, which is a comparable with other studies with relatively small sample sizes and is expected to climb close to $h^2_{SNP}$ with increasing sample sizes[7].

*2.3 Internal validation of lead SNPs*

We examined the replicability of the discovery lead SNPs as defined in FUMA in our holdout sample ($N_{FC}$ = 3,408; $N_{SC}$ = 3,412). From these six lead SNPs, we estimated to replicate three (exact 3.99) at $\alpha$ = (0.05/ n lead SNPs per trait) in our holdout sample given their winner's curse corrected effect size and the sample sizes of the discovery and replication samples. Observations showed three discovery lead SNPs to be significant (Supplementary Table 13). Figure S2 shows the probability distributions for all six discovery lead SNPs to be significant at increasing replication sample sizes.

### Supplementary Figures

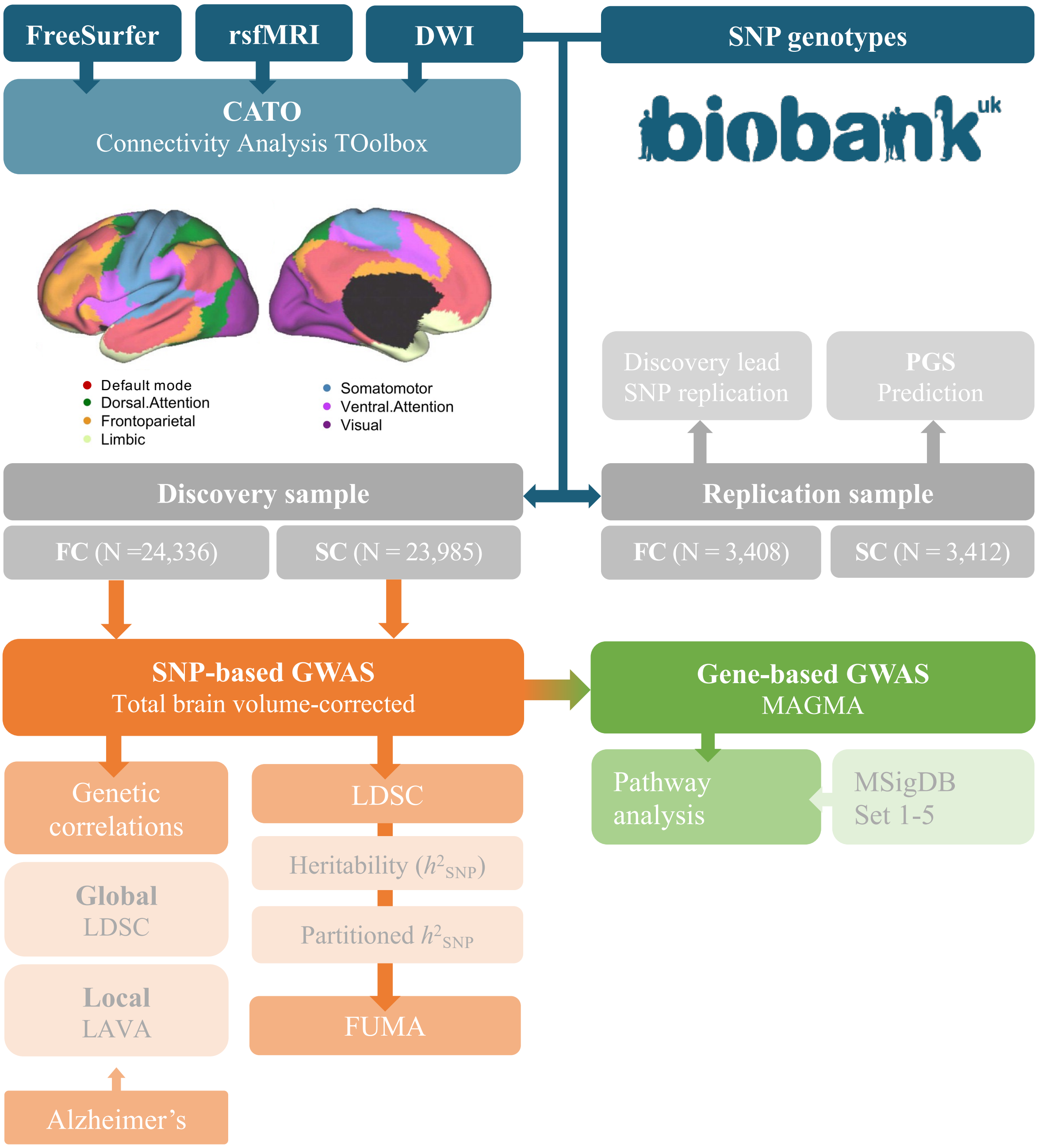


*Figure S1*. Flowchart of Methods involved in the current study. Functional and structural connectivity (FC/SC) within resting-state networks (RSN) as defined by Yeo et al[8] were obtained similarly as previously described[9]. SNP- and gene-based GWAS and in silico follow-up were performed on a discovery sample and were validated in a replication sample.

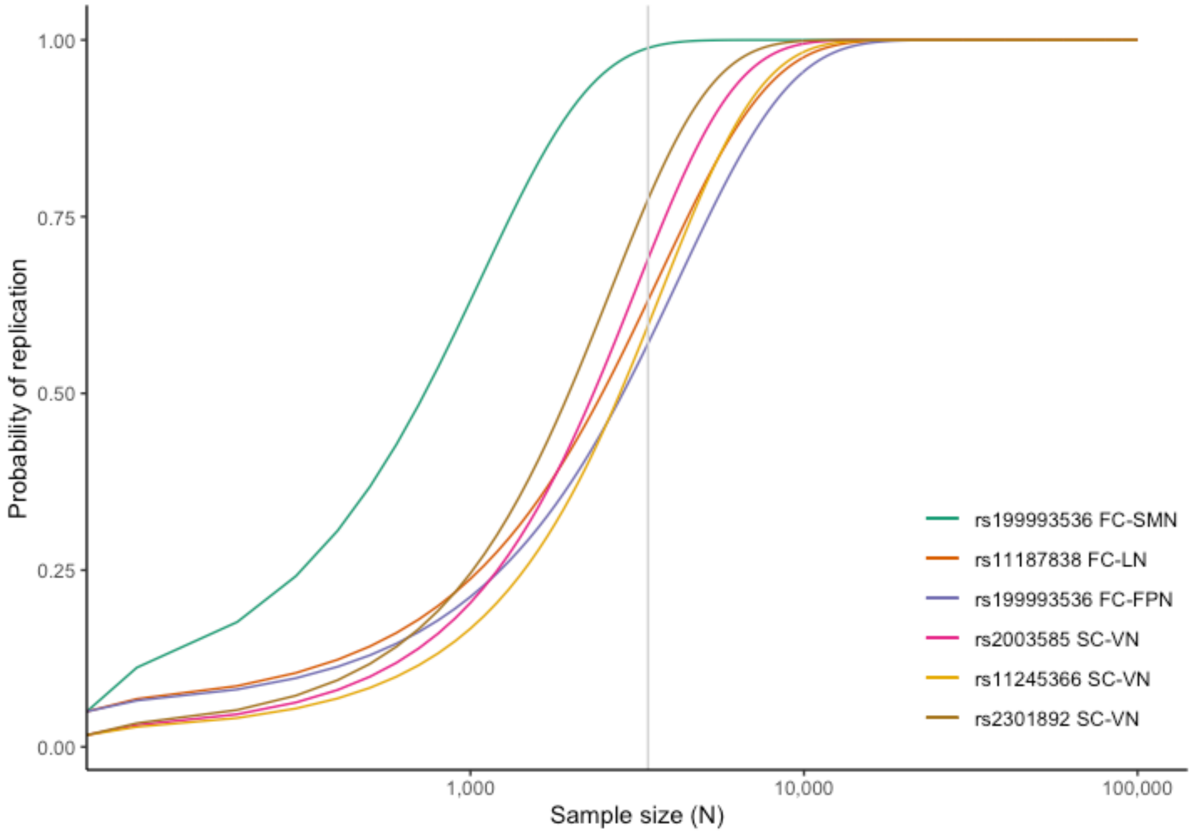


*Figure S2*. Probability curves of discovery lead SNPs being significant in a replication sample of increasing size (*log10*(N) scale). The current replication sample size of this study is represented by the vertical grey line. Please refer to Supplementary Methods section 1.4 for further details. Abbreviations: functional connectivity (FC), structural connectivity (SC), somatomotor network (SMN), limbic network (LN), frontoparietal network (FPN), visual network (VN).

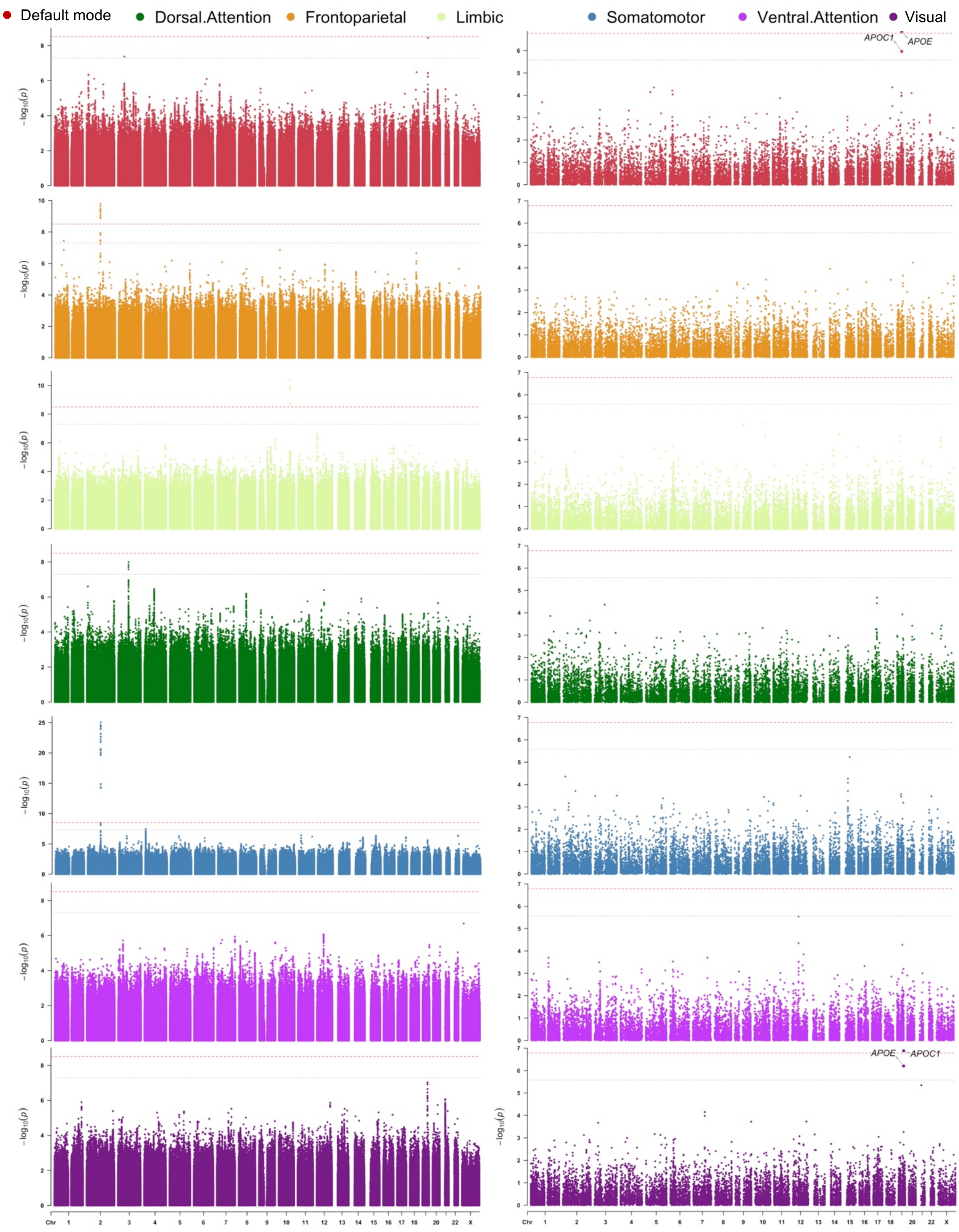


*Figure S3*. Manhattan plots of a) SNP-based and b) gene-based GWAS for RSN-FC. The light grey dashed horizontal line indicates a) traditional GWS ($p < 5\times10^{-8}$) or b) significance after correcting for the number of genes tested per trait ($p < 2.65\times10^{-6}$). The red dashed horizontal line indicates significance after an additional correction for the number of traits tested a) ($p < 3.13\times10^{-9}$) or b) ($p < 1.66\times10^{-7}$).

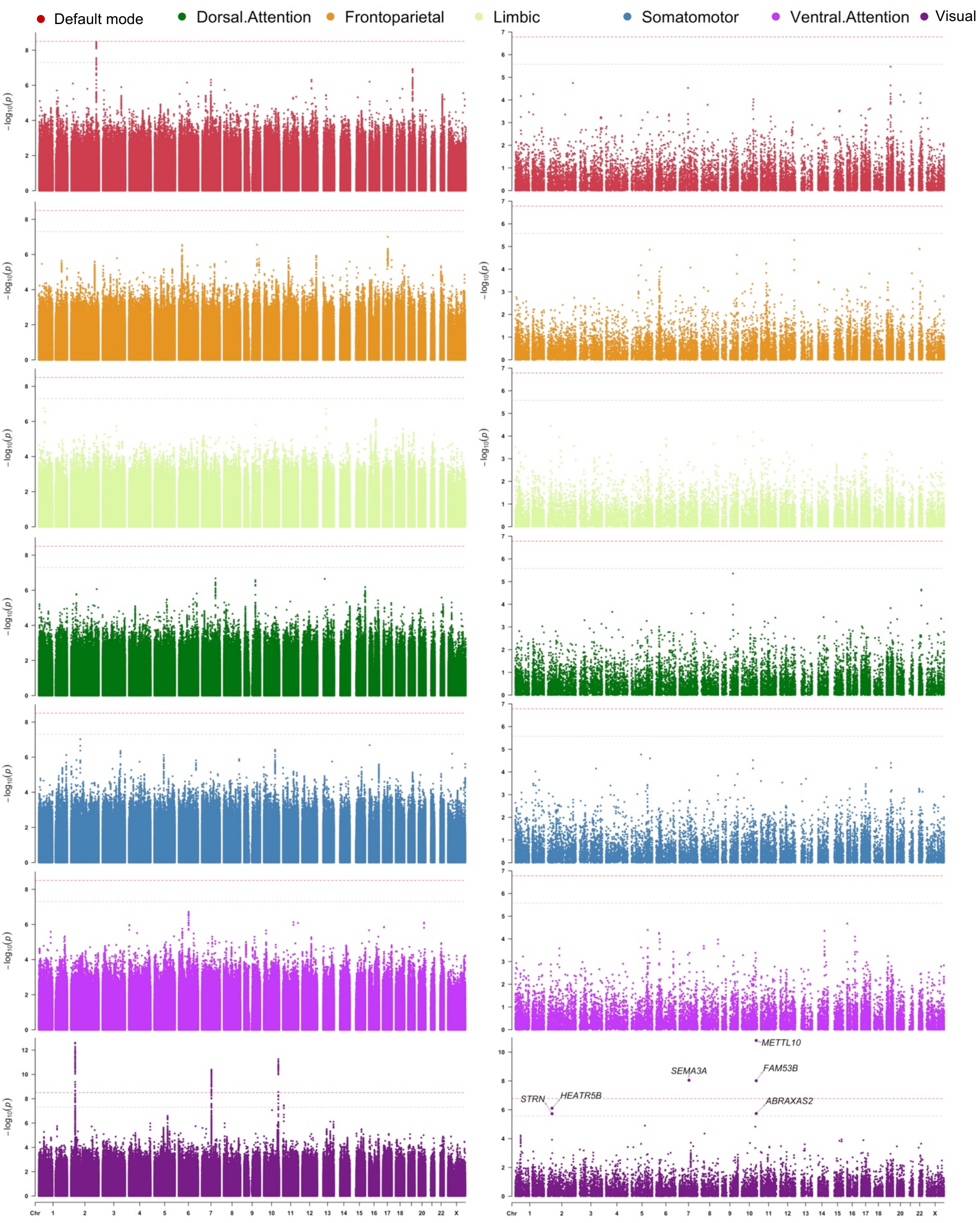


*Figure S4.* Manhattan plots of a) SNP-based and b) gene-based GWAS for RSN-SC. The light grey dashed horizontal line indicates a) traditional GWS ($p < 5\times10^{-8}$) or b) significance after correcting for the number of genes tested per trait ($p < 2.65\times10^{-6}$). The red dashed horizontal line indicates significance after an additional correction for the number of traits tested a) ($p < 3.13\times10^{-9}$) or b) ($p < 1.66\times10^{-7}$).

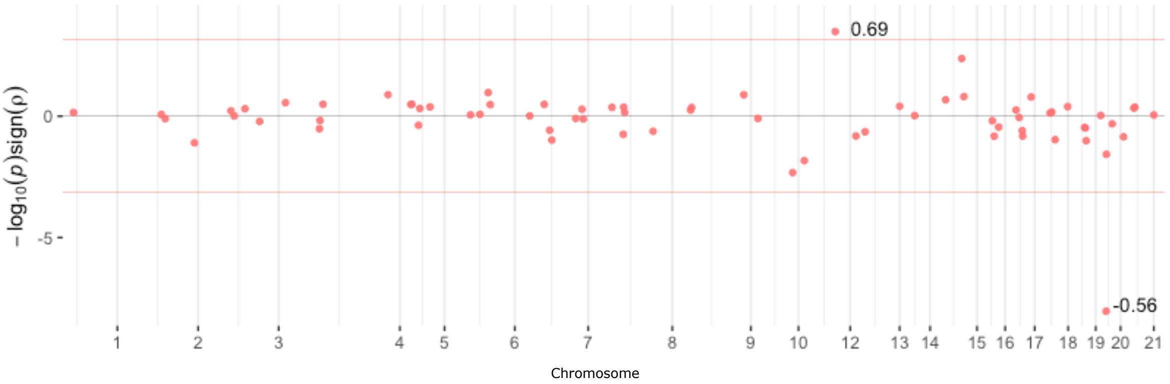


*Figure S5.* Local $r_g$ between default mode network-FC and Alzheimer's Disease as performed in LAVA. Only loci that passed the univariate $h^2$ threshold ($p < 1\times10^{-4}$) were tested for bivariate $r_g$, resulting in the Bonferroni-corrected significance threshold represented by the red line. Significant loci are visualized with their $r_g$ estimate. Within these loci, global FC did not show significant univariate $h^2$ and could therefore not bias these results. See Supplementary Table 7 for all local $r_g$ summary statistics.
